# Withdrawal from Chronic Ethanol Exposure Increases Postsynaptic Glutamate Function of Insular Cortex Projections to the Rat Basolateral Amygdala

**DOI:** 10.1101/772053

**Authors:** Molly M. McGinnis, Brian C. Parrish, Brian A. McCool

**Affiliations:** Department of Physiology and Pharmacology Wake Forest School of Medicine Winston-Salem, NC 27157

## Abstract

A key feature of alcohol use disorder (AUD) is negative affect during withdrawal, which often contributes to relapse and is thought to be caused by altered brain function, especially in circuits that are important mediators of emotional behaviors. Both the agranular insular cortex (AIC) and the basolateral amygdala (BLA) regulate emotions and are sensitive to ethanol-induced changes in synaptic plasticity. The AIC and BLA are reciprocally connected, however, and the effects of chronic ethanol exposure on this circuit have yet to be explored. Here, we use a combination of optogenetics and electrophysiology to examine the pre- and postsynaptic changes that occur to AIC – BLA synapses following withdrawal from 7- or 10-days of chronic intermittent ethanol (CIE) exposure. While CIE/withdrawal did not alter presynaptic glutamate release probably from AIC inputs, withdrawal from 10, but not 7, days of CIE increased AMPA receptor-mediated postsynaptic function at these synapses. Additionally, NMDA receptor-mediated currents evoked by electrical stimulation of the external capsule, which contains AIC afferents, were also increased during withdrawal. Notably, a single subanesthetic dose of ketamine administered at the onset of withdrawal prevented the withdrawal-induced increases in both AMPAR and NMDAR postsynaptic function. Ketamine also prevented the withdrawal-induced increases in anxiety-like behavior measured using the elevated zero maze. Together, these findings suggest that chronic ethanol exposure increases postsynaptic function within the AIC – BLA circuit and that ketamine can prevent ethanol withdrawal-induced alterations in synaptic plasticity and negative affect.

## Introduction

Alcohol use disorder (AUD) is characterized by prolonged and excessive alcohol consumption. This type of chronic alcohol exposure results in neuroadaptations in brain circuits that are crucial for the regulation of emotional states [1]. Alcohol withdrawal is associated with a negative-affective state, including an increase in anxiety, which is caused by disrupting the delicate balance between inhibitory and excitatory neurotransmission [2, 3]. For example, brain glutamate levels were significantly increased and GABA concentrations were significantly lower during acute withdrawal in alcohol-dependent patients as compared to healthy controls measured using magnetic resonance spectroscopy [4] or cerebral spinal fluid [5]. In preclinical models of dependence, chronic ethanol exposure and withdrawal produce pronounced increases in glutamatergic synaptic function in the basolateral amygdala (BLA), which likely contributes to increased anxiety-like behavior expressed during ethanol withdrawal [6]. The BLA serves as the primary input nuclei in the amygdala’s emotional-related neural circuitry [7], receiving highly processed sensory information from cortical afferents arriving via the external capsule (EC) [8]. Our laboratory has extensively characterized both the pre- and postsynaptic changes that occur in the BLA as a result of chronic intermittent ethanol (CIE) exposure and withdrawal (WD). For instance, we have reported using electrical stimulation that EC cortical glutamatergic inputs onto BLA principal neurons undergo predominantly postsynaptic alterations characterized by increased AMPA receptor function following CIE and WD [9, 10]. Although EC afferents onto BLA neurons potentially arise from many different cortical regions, it is unclear if CIE/WD-induced postsynaptic changes can be localized to individual circuits.

Tracing studies have demonstrated that the agranular insula cortex (AIC), which is located on the ventrolateral surface of the cerebral cortex rostrally and is a subdivision of the polymodal association cortex, is reciprocally connected with the BLA via the EC [11–14]. In addition to sensory inputs, the AIC integrates affective, anticipatory, and reward-related information coming from limbic regions [15]. Recent functional imaging studies in humans have identified the insula as a central region across many psychiatric and neurological disorders [16]. Importantly, many of the anatomical and functional features of the AIC are conserved across species, allowing the study of insular functions in animal models. A large body of evidence supports a role for the AIC in mediating fear and anxiety related behaviors as well as addiction. Specifically, pharmacological inactivation of the AIC decreases operant responding for alcohol along with a decrease in alcohol intake [17]. Several studies using opto- and chemogenetics have established a role of the AIC to nucleus accumbens (NAc) pathway in mediating alcohol intake in animal models of AUD. For example, optogenetic inhibition of glutamatergic AIC inputs to the nucleus accumbens reduces quinine-resistant alcohol intake, suggesting this pathway sustains aversion-resistant alcohol intake [18]. Additionally, chemogenetic silencing of AIC – NAc projections decreases alcohol intake in rats trained to self-administer alcohol [19]. Despite mounting evidence that the AIC is involved in AUD-related behaviors, very few studies have examined ethanol modulation of synaptic transmission and plasticity in the AIC. One study examining the acute effects of ethanol on GABA and glutamate transmission in the AIC found that glutamate, but not GABA, systems are sensitive to pharmacologically relevant concentrations of ethanol [20]. Specifically, ethanol had no effect on spontaneous inhibitory postsynaptic currents mediated by GABA_A_ receptors. However, ethanol inhibited N-methyl-D-aspartate receptor (NMDAR)-mediated excitatory postsynaptic currents in a concentration-dependent manner and acute, intoxicating concentrations of ethanol prevented the ability to induce LTD in the AIC [20]. However, the effects of chronic ethanol exposure on AIC circuits, such as AIC – BLA projections, are yet to be examined.

Since AIC inputs arrive at the BLA via the EC, and our laboratory has previously shown that EC – BLA synapses undergo postsynaptic changes following withdrawal from chronic ethanol exposure, we hypothesized that AIC – BLA synapses would express a similar form of ethanol-induced plasticity. To test this hypothesis, we exposed rats to CIE using ethanol vapor chambers, which is a commonly used method for inducing dependence in rodent models of AUD [21]. We’ve previously reported that varying lengths of this type of ethanol exposure produces behavioral alteration indicative of a dependence-like phenotype including increased voluntary ethanol-self administration and increased anxiety-like behavior during withdrawal [9, 22]. Using a combination of optogenetics and electrophysiology to record optically-evoked glutamatergic response in the BLA principal neurons that were innervated by Channelrhodopsin-expressing AIC terminals, we characterized the presynaptic and postsynaptic functions of AIC – BLA synapses in CIE- and air-exposed animals.

## Materials and Methods

### Animals

Male Sprague-Dawley rats were purchased from Envigo (Indianapolis, IN) and given access to food and water *ad libitum* upon arrival. Rats were pair-housed in a humidity and temperature-controlled room and maintained on a reverse 12:12h light-dark cycle (lights off at 9 AM). Rats that underwent surgery (N=96) were aged ∼5 weeks (100g) at arrival and rats that did not undergo surgery (N=32) were aged ∼8 weeks (250g) at arrival. All rats were aged ∼10 weeks (300g) at the time of behavioral manipulations and electrophysiology recordings. All animal care procedures were in accordance with the NIH Guide for the Care and Use of Laboratory Animals. All experimental procedures were approved in advance by the Institutional Animal Care and Use Committee at Wake Forest University Health Sciences.

### Stereotaxic Surgery

Rats were kept under continuous isoflurane anesthesia (3-5% for induction, 1-3% for maintenance) with oxygen flow at 1 L/min throughout the surgery. An adeno-associated viral vector containing Channelrhodopin (AAV5-CamKIIα-hChR2(H134R)-EYFP; UNC Vector Core, Chapel Hill, NC) was bilaterally microinjected (1 μL/side) into the agranular insular cortex (AIC) using a Neurostar StereoDrive (Germany) with the following coordinates relative to bregma (in mm): 2.76 AP, ± 3.50 ML, 5.10 DV. The virus was delivered at a rate of 0.1 μL/min over 10 min using a Harvard Apparatus pump (Holliston, MA). Injectors were left in place for an additional 5 min after injection to allow for virus to diffuse. Rats were given 2 mL of warmed sterile saline and 3 mg/kg ketoprofen (Ketofen; Patterson Veterinary, Devens, MA) for pain management at the end of the surgery. Sutures were removed and rats were pair-housed 1 week following surgery. A total of 4 weeks was allowed for the rats to recover and for virus expression prior to experimentation. Injection sites were confirmed by visualizing EYFP in coronal slices of the AIC using fluorescence microscopy post-mortem. If there was unintended viral spread, rats were excluded.

### Chronic Intermittent Ethanol (CIE) Vapor Exposure

Using standard procedures from our laboratory [9], rats were exposed to CIE for 7 or 10 consecutive days. Pair-housed rats in their home cages were placed inside larger, custom-build Plexiglas chambers (Triad Plastics, Winston-Salem, NC). Ethanol vapor was pumped into the chambers beginning at 9 PM (start of the light cycle) at a constant rate of 16 L/min and maintained at ∼25 mg/L throughout the exposure for 12 h/day. Control animals were only exposed to room-air. All rats were weighed daily. Blood ethanol concentrations (BECs) were determined by a standard, commercially available alcohol dehydrogenase/NADH enzymatic assay (Carolina Liquid Chemistries, Greensboro, NC). Tail blood samples were collected periodically throughout the CIE exposure to monitor and adjust ethanol vapor levels as necessary. The standard BEC range that we’ve found to produce dependence in our laboratory is between 150-275 mg/dL. Average BECs in the CIE animals were 259.40 ± 9.26 mg/dL, which is within our range. All behavioral experiments and electrophysiology recordings were conducted 24 h after the last ethanol or air exposure.

### Drugs

Ketamine HCl (KetaVed; Patterson Veterinary, Devens, MA) was diluted in saline and administered at 10 mg/kg, IP. Rats received either ketamine injections or saline (control) injections at the onset of withdrawal, 24 h before behavioral testing and electrophysiology recording.

### Elevated Zero Maze

Rats were tested on the elevated zero maze (EZM; Med Associates, Fairfax, VT) 24 h after the last ethanol or air exposure to assess anxiety-like behavior. In experiments were ketamine or saline administered, injections occurred 24 h prior. The circular EZM consists of two open sections that are dimly lit (∼40 lux) and two dark enclosed sections. In order to assess anxiety-like behavior and general locomotion the center point, tail base, and nose point of the rats were tracked throughout the 5 min test using a Basler ace monochrome camera (Basler AG, Germany) and EthoVision XT video tracking software (Noldus; Leesburg, VA). The apparatus was cleaned with warm water and mild soap and then thoroughly dried between animals. Data are represented as percentage of time, which was calculated by dividing the total amount of time spent in both open areas by the entire duration of the test.

### Electrophysiology

#### Slice Preparation

Rats were deeply anesthetized with isoflurane and decapitated with a guillotine. Brains were quickly removed and incubated in ice-cold sucrose-modified artificial cerebral spinal fluid (aCSF) for 5 min. The sucrose aCSF contained (in mM): 180 Sucrose, 30 NaCl, 4.5 KCl, 1 MgCl_2_·6H_2_O, 26 NaHCO_3_, 1.2 NaH_2_PO_4_, 10 D-glucose, 0.10 ketamine and was equilibrated with 95% O_2_ and 5% CO_2_. 400-micron thick coronal slices containing the BLA were collected using a VT1200/S vibrating blade microtome (Leica, Buffalo Grove, IL) and incubated at room temperature (∼25°C) in oxygenated standard aCSF for ≥ 1h prior to recordings. The standard aCSF solution contained (in mM): 126 NaCl, 3 KCl, 1.25 NaH_2_PO_4_, 2 MgSO_4_·7H_2_O, 26 NaHCO_3_, 10 D-glucose, and 2 CaCl_2_·2H_2_O. Unless otherwise noted, all chemicals were obtained from Tocris (Ellisville, Missouri) or Sigma-Aldrich (St. Louis, MO).

#### Whole-Cell Patch-Clamp Recording

Using standard methods for whole-cell voltage-clamp electrophysiology from our laboratory [9], we recorded synaptic responses from BLA slices kept in a submersion-type recording chamber that was continuously perfused with oxygenated, room temperature (∼25°C) aCSF at a rate of 2 mL/min. Recordings electrodes were filled with an intracellular solution containing (in mM): 145 CsOH, 10 EGTA, 5 NaCl, 1 MgCl2·6H2O, 10 HEPES, 4 Mg-ATP, 0.4 Na-GTP, 0.4 QX314, 1 CaCl_2_·2H_2_O. The osmolarity was adjusted to ∼285 Osm/L using sucrose and pH was adjusted to ∼7.3 using gluconic acid. Synaptic glutamate currents were recorded at a membrane holding potential of -65 mV and pharmacologically isolated using the GABA_A_ antagonist, picrotoxin (100 μM). For NMDA-mediated EPSCs, extracellular Mg^2+^ concentrations were lowered to 0.2mM. Data were acquired using a Axopatch 700B amplifier (Molecular Devices, Foster City, CA) and were low-pass filtered at 2 kHz. pClamp 10 software (Molecular Devises, Foster City, CA) was used for later analysis. BLA principal neurons were included based on their electrophysiological characteristics of low access resistance (≤ 25 MΩ) and high membrane capacitance (> 100 pF) [23]. Cells in which capacitance or access resistance changed ≥ 20% during the recording or that did not meet principal neuron criteria were excluded from analysis.

#### Optogenetics

Optogenetic responses were evoked using a 473 nm laser connected to a fiber optic cable (Thorlabs, Newton, NJ). The naked end of the cable was placed just above the external capsule on the lateral side of the BLA. Five-millisecond laser pulses were used to activate Channelrhodopsin found in the AIC terminals. Sweeps were recorded every 30 sec and light stimulation intensities were submaximal and normalized to elicit synaptic responses with amplitudes ∼100 pA. In some recordings, 1 μM tetrodotoxin (TTX; Tocris) and 20 mM 4-aminopyridine (4-AP; Tocris) were included for more stringent isolation of monosynaptic transmission [24, 25].

#### Paired-Pulse Ratio

Two 5msec light stimuli of equal intensity were delivered to the external capsule at an inter-stimulus interval of 50 msec. This short interval is traditionally viewed as an indicator of presynaptic release probability [26]. The paired-pulse ratio (PPR) was conservatively calculated using the evoked EPSC amplitudes as: ([Peak 2 amplitude – Peak 1 amplitude]/Peak 1 amplitude). The average paired-pulse ratio was determined from a 5-min, 11 sweep recording.

#### Strontium Substitution

As a measure of input-specific postsynaptic function, a strontium (Sr^2+^) substitution method was used whereby extracellular calcium (2mM) was replaced by strontium (2mM) in the bath aCSF. This technique allows for the measurement of asynchronous excitatory postsynaptic potentials (aEPSCs) [27], with changes in frequency providing a measure of calcium-independent presynaptic function and amplitude providing a measure of postsynaptic efficacy. As described previously by our laboratory [10], a bipolar stimulating electrode or an optical fiber was placed at the external capsule where an electrical or light stimulation was applied every 30 sec. Semi-automated aEPSC analysis was conducted on responses starting 50 msec post-stimulation to include responses during a 400 msec window. The median inter-event interval time and amplitude of individual aEPSC events were extracted using the Mini Analysis Program (Synaptosoft, Fort Lee, NJ).

#### NMDA Input – Output

To record NMDA-mediated EPSCs, neurons were voltage-clamped at - 60 mV holding potential in the presence of low extracellular magnesium (0.2 mM), picrotoxin (100 μM), and the AMPA receptor antagonist, 6,7-dinitroquinoxaline-2,3-dione (DNQX; 20 μM). EPSCs were electrically evoked every 30 sec by brief (0.2 msec) square-wave stimulations delivered to the external capsule using platinum/iridium concentric bipolar stimulating electrodes (FHC, Bowdoinham, ME) with an inner pole diameter of 12.5 μm. Increasing stimulus intensities (10 μA to 200 μA) produced graded monosynaptic responses.

### Statistics

All data are represented as mean ± SEM throughout the text and figures. Primary statistical analyses were conducted using Prism 5 (GraphPad, La Jolla, CA). Differences between groups were analyzed using t-tests and one- or two-way ANOVA, depending on the experimental design. A value of *p* < 0.05 was considered statistically significant. When significant effects were obtained using ANOVA tests, Bonferroni post hoc comparisons between groups were performed.

## Results

### Neurons in the AIC make monosynaptic glutamatergic synapses onto principal neurons in the BLA

Using anterograde and retrograde tracing, previous studies have shown that the AI sends projections to the BLA [12, 14]. Microinjection of Channelrhodopsin-eYFP into the AIC (Fig. 1A), encompassing the dorsal and ventral agranular insular areas, resulted in dense eYFP fluorescence in the BLA (Fig. 1B). By recording from principal neurons in the BLA and optically stimulating ChR2-expressing AIC terminals (Fig. 1C), we established baseline EPSCs and then washed on the sodium channel blocker, TTX (1 μM), which abolished the light-evoked EPSC. With the addition of the potassium channel inhibitor 4-AP (20 mM), we were able to rescue the light-evoked EPSCs (Fig. 1D). On average, the peak amplitude of EPSCs recorded in the presence of TTX and 4-AP were 81.38% ± 2.008 of the baseline peak amplitude, demonstrating the vast majority of light-evoked responses were monosynaptic. Importantly, addition of the AMPA receptor antagonist, DNQX (20 uM), completely abolished the monosynaptic light-evoked ESPC, suggesting the responses were glutamatergic. Thus, AIC – BLA synapses onto principal are monosynaptic and glutamatergic.

**Figure 1.**
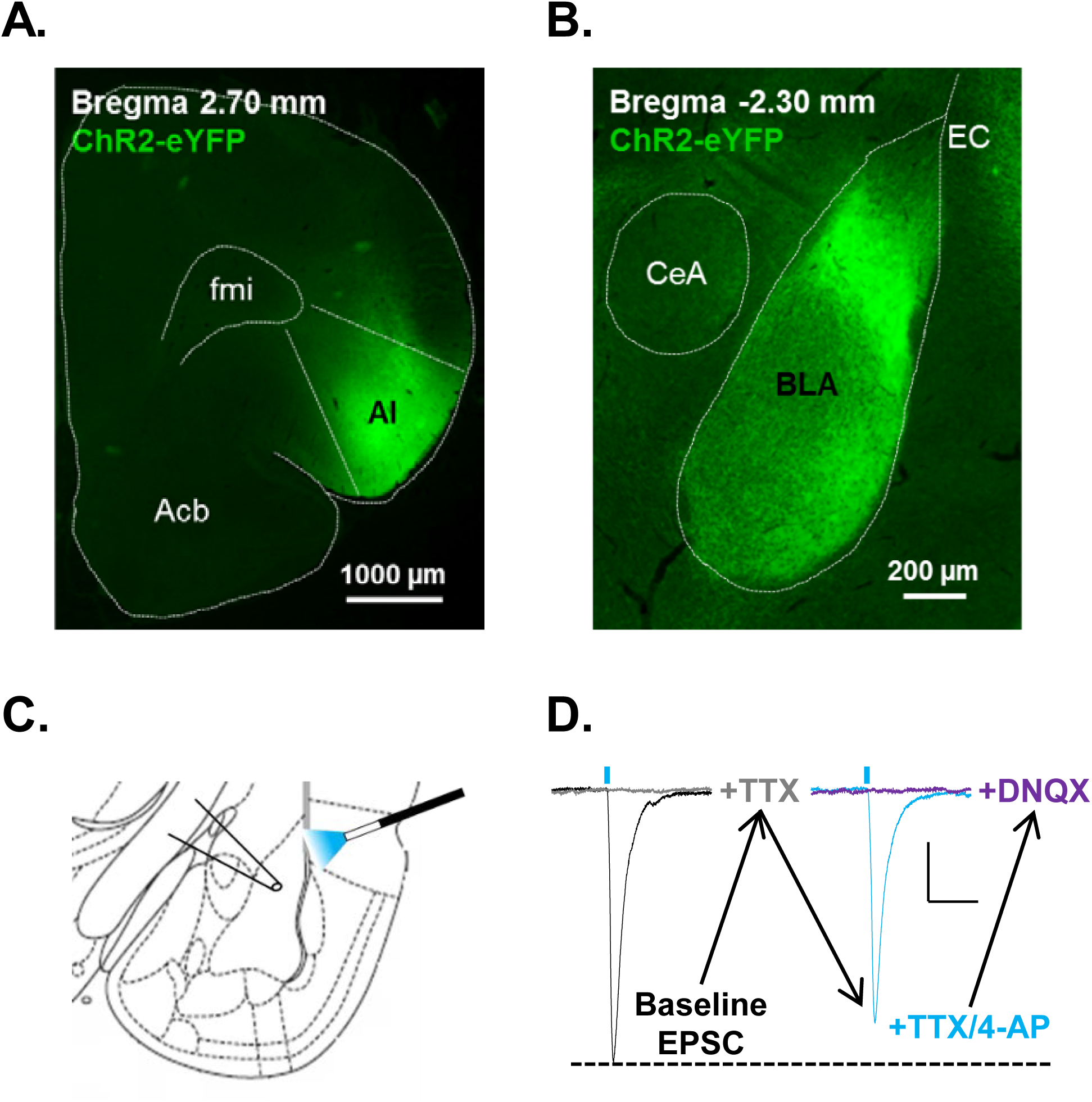
The agranular insular cortex sends monosynaptic glutamatergic projections to basolateral amygdala principal neurons. **A**, Representative fluorescent image of the agranular insular cortex (AI) injection site and **B**, resulting terminal field in the basolateral amygdala (BLA) 4 weeks after injection of Channelrhodopsin. **C**, Schematic depicting the typical placement of the optical fiber and patch electrode for electrophysiology recordings. The optical fiber used to deliver 470 nm blue light was placed just outside of the BLA above the lateral external capsule (EC) to activate the Channelrhodopsin-expressing terminals entering the BLA from the AI. BLA principal neurons were patched with recording electrodes that were placed near the stimulating fiber where the YFP-expressing terminals were most dense. **D**, Representative traces of optogenetically-evoked EPSCs recorded from AI – BLA synapses at baseline (with picrotoxin, 100μM; black trace) and in the presence of tetrodotoxin (TTX, 1μM; gray trace), 4-aminopyradine (4-AP, 20mM; cyan trace), and 6,7-dinitroquinoxaline-2,3-dione (DNQX, 20μM; purple trace). Blue dashes represent optogenetic stimulation (5msec) with blue light. Scale bars= 25pA X 50msec. Acb= nucleus accumbens, AI= agranular insular cortex, BLA= basolateral amygdala, CeA= central nucleus of the amygdala, EC= external capsule, fmi= anterior forceps of corpus callosum.

### AIC – BLA synapses do not undergo presynaptic changes during withdrawal from chronic ethanol exposure

Using electrical stimulation, several studies from our laboratory have reported a lack of treatment effect on presynaptic function at glutamatergic inputs arriving at BLA principal neurons via the external capsule [9, 10]. However, in a recent study using periadolescent rats, we reported post-synaptic facilitation of these inputs after 7 days of CIE [9]. To examine whether AIC – BLA synapses express either pre- or post-synaptic effects of chronic ethanol, we exposed animals to 7 or 10 consecutive days of CIE (Fig. 2A) and recorded PPRs from principal neurons in the BLA 24h into withdrawal by optically stimulating AIC terminals arriving via the external capsule. We found no difference in PPRs recorded from AIC – BLA synapses in AIR (PPR = -0.19 ± 0.06, N=15) and 7d CIE/WD (-0.17 ± 0.08, N=15) animals (Fig. 2B; Unpaired t-test, t(28)= 0.245, p= 0.809). Similarly, after exposing animals to a longer 10 day CIE exposure which produces dependence-like physiological and behavioral alterations [9, 10, 22, 28–31], we again found no difference in PPRs recorded from AIC – BLA synapses in AIR (PPR= -0.19 ± 0.05, N=15) and 10d CIE/WD (PPR= -0.15 ± 0.06, N=15) animals (Fig. 2C; Unpaired t-test, t(28)= 0.406, p= 0.688).

**Figure 2.**
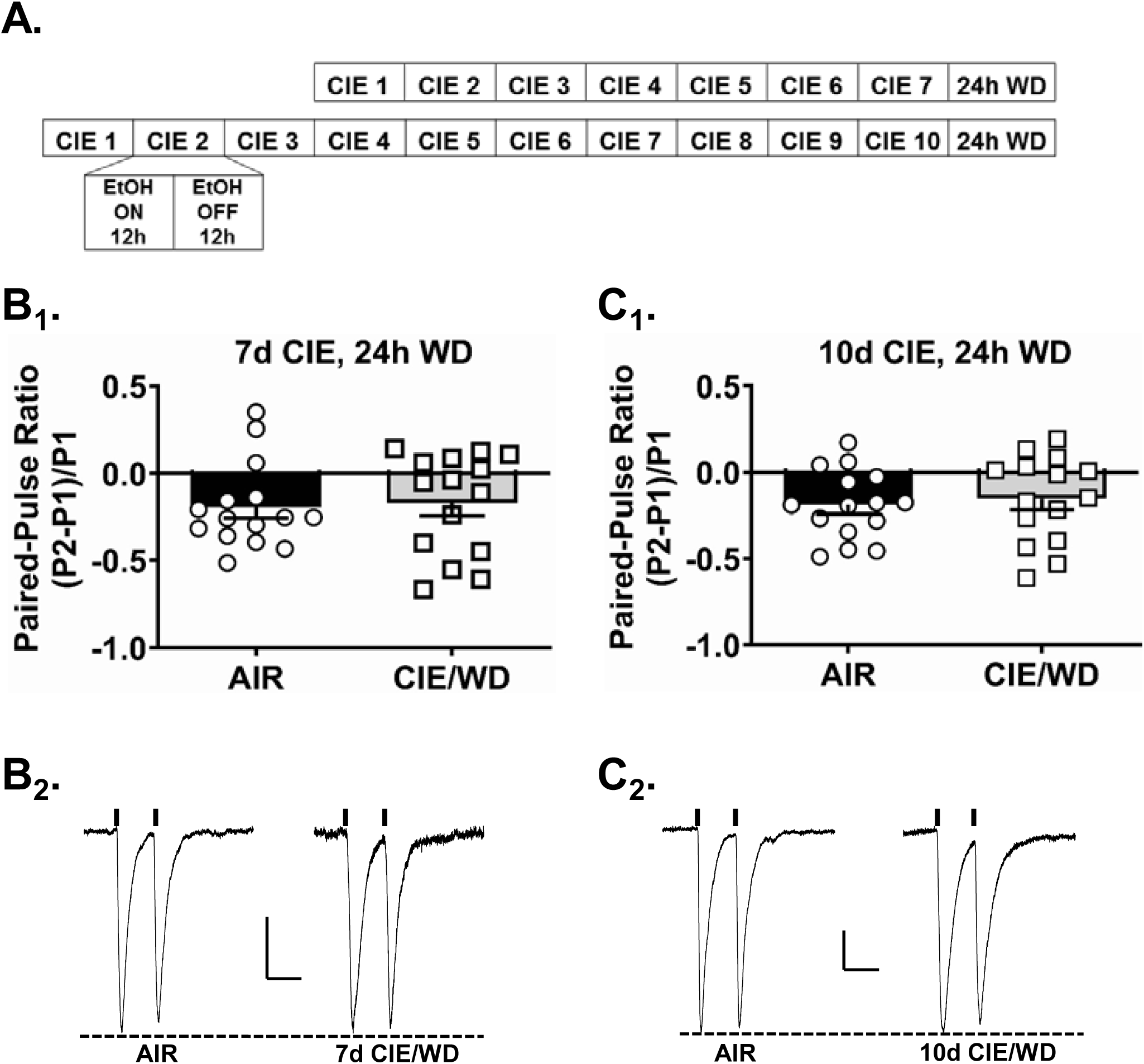
Withdrawal from chronic ethanol exposure does not alter presynaptic glutamate release from insular terminals onto basolateral amygdala principal neurons. **A**, Experimental timeline depicting the 7- or 10- day chronic intermittent ethanol (CIE) exposures. Each day, rats were exposed to 12-hours of ethanol (EtOH) and 12-hours of air. All electrophysiology recordings were conducted 24-hours after the last ethanol exposure. Control animals were identically housed but only exposed to air. **B_1_**, Optogenetically-evoked glutamate paired-pulse ratios (PPRs, 50msec inter-stimulus interval) recorded from AI – BLA synapses after 7 days of CIE were not different between AIR (N= 15) and CIE/WD (N= 15) animals. **C_1_**, PPRs recorded from AI – BLA synapses after 10 days of CIE were not different between AIR (N= 15) and CIE/WD (N= 15) animals. **B_2_**, Representative traces of PPRs recorded from AIR (left) and CIE/WD (right) after 7- and **C_2_**, 10-days of CIE. Unpaired t-test, p > 0.05. Vertical lines about the traces indicate timing of the light pulse. Scale bars= 20pA X 50msec.

### Withdrawal from 10, but not 7, days of CIE increases postsynaptic function at AIC – BLA synapses

By substituting strontium for calcium in the bath aCSF, we measured the frequency and amplitude of light-evoked, input-specific, AMPA-mediated aEPSCs at AIC – BLA synapses 24h into withdrawal after 7 or 10 days of CIE (or air) exposure. After 7 days of CIE, we found no change in the aEPSC frequency (Fig. 3A_1_; Unpaired t-test, t(16)= 0.939, p= 0.362) between AIR (13.61 ± 0.28 msec, N=10) and 7d CIE/WD animals (13.20 ± 0.34 msec, N=8). There was also no change in the aEPSC amplitude (Fig. 3A_2_; Unpaired t-test, t(16)= 1.035, p=0.316) between AIR (8.68 ± 0.15pA) and 7d CIE/WD animals (8.36 ± 0.29pA, N=8). After 10 days of CIE, we again found no change in the aEPSC frequency (Fig. 3B_1_; Unpaired t-test, t(32)= 1.008, p=0.321) between AIR (13.63 ± 0.27msec, N=17) and 10d CIE/WD animals (14.11 ± 0.39, N=17). However, there was a significant increase in the aEPSC amplitude following 10d CIE/WD (Fig. 3B_2_; Unpaired t-test, t(32)= 10.94, p< 0.0001; AIR = 8.06 ± 0.18pA, CIE/WD = 10.98 ± 0.20pA). These data suggest that 10d CIE/WD increases the postsynaptic function at AIC-BLA synapses.

**Figure 3.**
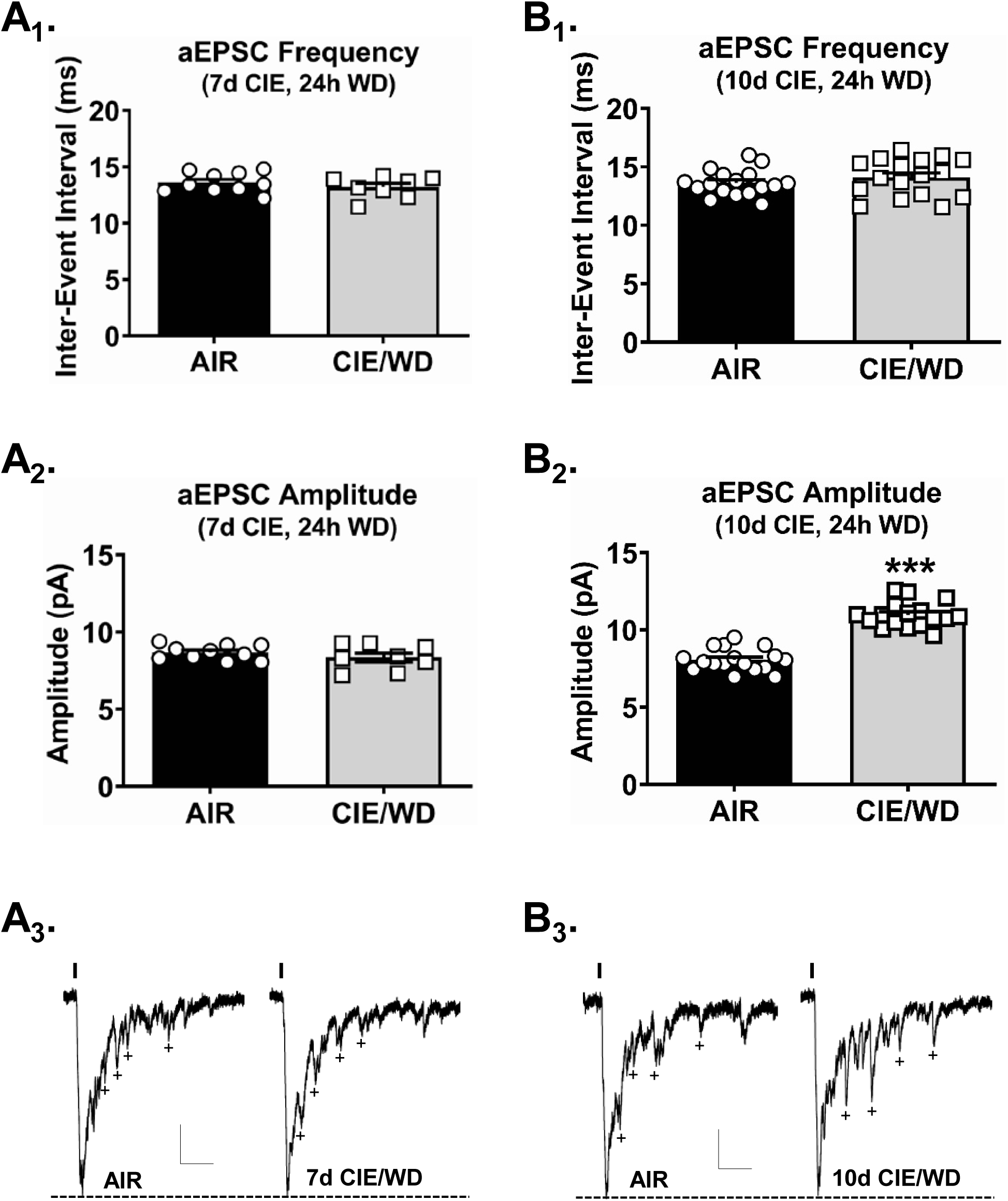
Withdrawal from 10, but not 7, days of chronic exposure enhances postsynaptic function at agranular insular synapses in the basolateral amygdala. **A**, The frequency (A_1_) and amplitude (A_2_) of optogenetically-evoked asynchronous excitatory postsynaptic currents (aEPSCs) were similar in AIR (N= 10) and 7d CIE/WD (N= 8) recordings from AI – BLA synapses. **B_1_**, Following 10d CIE/WD, there was no change in the frequency of aEPSC recorded from AIR (N= 17) and 10d CIE/WD (N= 17) animals. **B_2_**, The aEPSC amplitudes were significantly larger in 10d CIE/WD animals as compared to AIR. **A_3_**, Representative aEPSC traces from 7- and **B_3_**, 10-day CIE animals. Vertical lines indicate the position of the light pulse. ‘+’ represent exemplar aEPSCs used in the analysis. Scale bars= 30pA X 50msec. Unpaired t-test, ***p < 0.0001.

### Ketamine administration at the onset of withdrawal prevents the CIE/WD-induced increases postsynaptic function

It is well established that NMDA receptors play an integral role in activity-dependent long-term plasticity, such as LTP and LTD, that is expressed as changes in AMPA receptor-mediated transmission [32]. Our laboratory has previously reported that NMDARs in the BLA are sensitive to chronic ethanol [33] and withdrawal [28]. Further, recent work [34] showed that a single ketamine exposure at the onset of forced abstinence from long-term ethanol consumption can increase the capacity for NMDA-dependent LTP in the BNST. We therefore administered ketamine at the onset of withdrawal from CIE and examined AIC-BLA postsynaptic NMDA receptor function. Using electrical stimulation delivered to the external capsule, we recorded NMDAR-mediated EPSCs from BLA principal neurons across a range of stimulation intensities (Fig. 4C) in AIR and 10d CIE/WD animals that received either ketamine (10mg/kg) or saline (Fig. 4A). Mixed effects analysis found a significant interaction between stimulation intensity and treatment group (Fig. 4B; F(12, 156)=8.918, p<0.0001).

**Figure 4.**
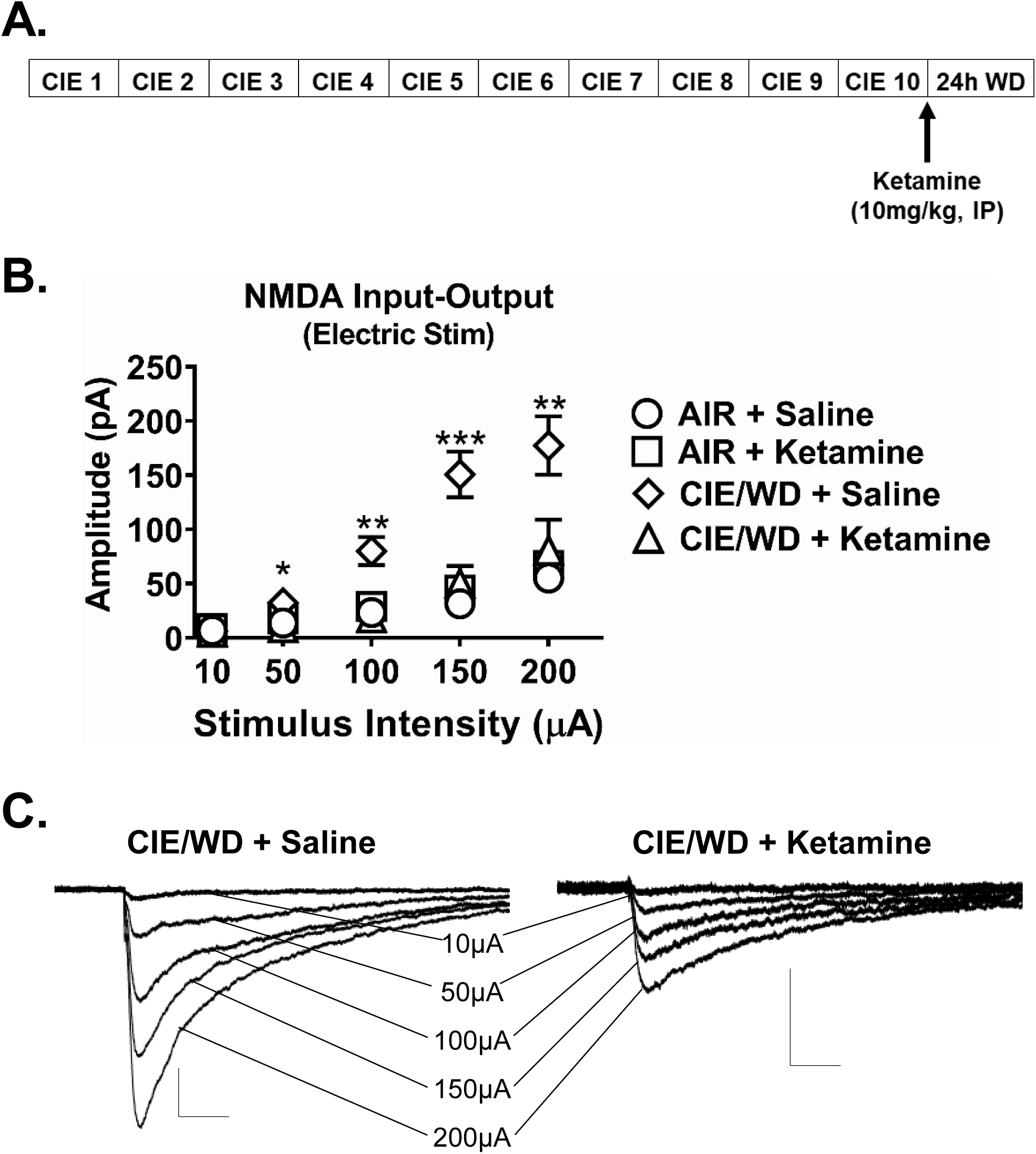
Ketamine administration at the onset of withdrawal prevents ethanol-induced increase in NMDA receptor function. **A**, Experimental timeline depicting the 10-day CIE exposure and ketamine injection at the onset of withdrawal, 24-hours prior to the electrophysiology recordings. **B**, Significant increase in NMDAR function in CIE/WD + Saline (N= 18) group as compared to AIR + saline (N= 14) group across multiple stimulation intensities (Two-way ANOVA, interaction, F(12, 156)=8.92, p<0.001; Bonferroni posttest, * - p<0.05, ** - p<0.01). There were no significant differences between AIR + saline, Air + ketamine (N= 10), and CIE/WD + ketamine (N= 9; all p>0.05, see text for details). **C**, Representative traces of electrically-evoked NMDAR-mediated EPSCs recorded from CIE/WD + Saline (left) and CIE/WD + Ketmaine (right) EC – BLA synapses. Scale bars= 50pA X 100msec.

Comparing treatment groups to the Air + Saline controls, Dunnett’s multiple comparison test indicated that only the CIE + Saline group was significantly different from control at the 50μA (p<0.05), 100μA (p<0.01), 150μA (p<0.001), and 200μA (p<0.01) stimulation intensities. These results demonstrate that NMDAR function at EC -BLA synapses is increased following 24h withdrawal from 10 days of CIE, and that administration of ketamine at the onset of withdrawal can block this effect.

After findings that ketamine administration could prevent the 10d CIE/WD-induced increase in NMDAR function at EC – BLA synapses (Fig. 4), we next examined the effect of ketamine treatment on postsynaptic AMPAR function at AIC – BLA synapses. Using animals injected with ChR2 in the AIC, we exposed them to 10 days of CIE (or air) and injected them with saline or ketamine at the onset of withdrawal and recorded light-evoked, input-specific aEPSCs from BLA principal neurons 24h later (Fig 5A). A two-way ANOVA on median aEPSC frequency (Fig. 5A_1_) revealed no main effect of exposure (AIR vs CIE/WD; F(1,43)= 2.857, p>0.05), no main effect of treatment (Saline vs Ketamine; F(1,43)= 0.2152, p>0.05), and no interaction (F(1,43)= 0.5556, p>0.05). However, a two-way ANOVA on median aEPSC amplitude (Fig. 5A_2_) revealed a significant interaction between exposure and treatment (F(1,43)= 22.53, p<0.0001) as well as significant main effects (exposure, F(1,43)= 33.59, p< 0.0001; treatment, (F(1,43)= 28.19, p< 0.0001) for light-evoked aEPSCs. Bonferroni posttests indicated a significant (p < 0.001) increase in aEPSC amplitude in saline injected CIE/WD neurons (12.4 ± 0.6pA, N=13) compared to saline injected AIR-exposed controls (8.0 ± 0.3pA, N=10, t=7.35), ketamine-injected Air controls (8.0 ± 0.3pA, N=13, t=8.13), and ketamine-injected CIE/WD neurons (8.6 ± 0.5pA, N=13, t=7.36). There were no significant differences between ketamine injected CIE/WD neurons and either the Air-Saline (t=0.33) or Air-ketamine groups (t=0.75). These findings demonstrate that ketamine prevents the increase in AMPA-mediated aEPSC amplitude at AIC – BLA synapses.

**Figure 5.**
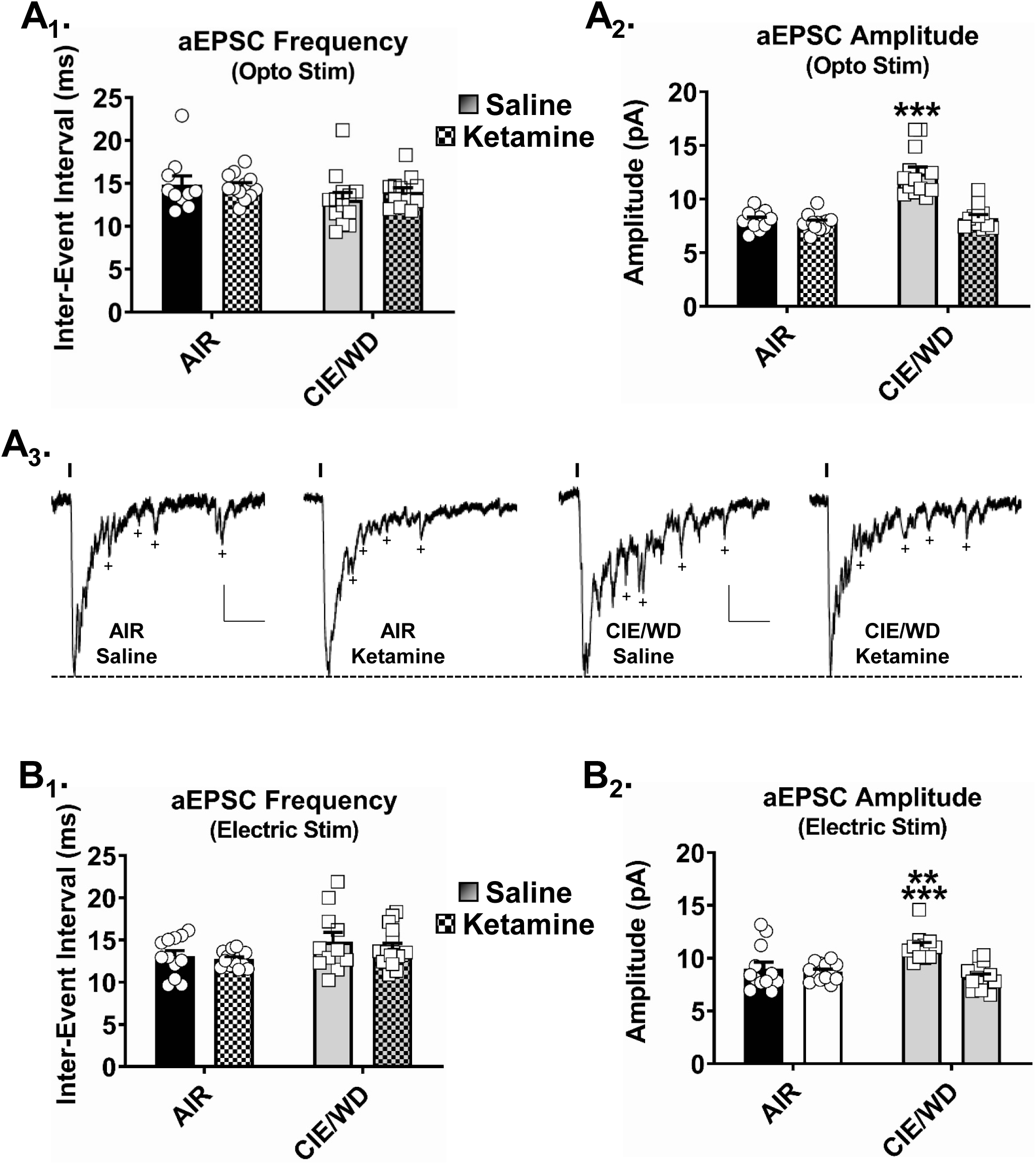
Ketamine administration at the onset of withdrawal attenuates CIE-induced increases in AMPA receptor function measured from both optically-stimulated AI – BLA synapses and electrically-stimulated EC – BLA synapses. **A_1_**, Neither CIE/WD nor ketamine changed the frequency of optically-evoked asynchronous excitatory postsynaptic currents (aEPSCs) measured from AI – BLA synapses. **A_2_**, While CIE/WD + Saline (N=12) increased aEPSC amplitude compared to AIR + Saline (N= 12), ketamine administration (10mg/kg, I.P., N=16) blocked this effect. Ketamine administration did not alter optically-evoked aEPSC amplitude in neurons (N=13) from air-exposed animals. Two-way ANOVA, interaction p<0.001 (F(1,43)=22.5), *** - p<0.001 with Bonferroni posttest CIE/WD + Saline versus cells from the other three treatment groups. **A_3_**, Representative traces of optogenetically-evoked responses from AI – BLA synapses. Scale bars= 30pA X 50msec. Examples of aEPSCs analyzed are marked (+). **B_1_**, Neither CIE/WD nor ketamine altered aEPSC inter-event intervals recorded from electrically-evoked EC-BLA responses. **B_2_**, Electically-evoked aEPSC amplitudes from EC-BLA responses were significantly greater (Two-way ANOVA, interaction p<0.01, F(1,48)=10.3) in CIE/WD+Saline neurons (N= 11) compared to neurons from Air+Saline (N=12, ** -- p<0.01, Bonferroni posttest), Air + ketamine (N= 13, ** -- p<0.01), and CIE/WD + ketamine (N = 16, *** - p<0.001).

Next, we wanted to determine if our findings with optogenetic stimulation of AIC – BLA synapses were generalizable to all EC – BLA inputs by using electrical stimulation. A two-way ANOVA on median frequency (Fig. 5B_1_) revealed no interaction (F(1,48)= 0.01, p>0.05) between the main factors of exposure (Air vs CIE/WD) and treatment (Saline vs Ketamine), no main effect of exposure (F(1,48)= 3.09, p>0.05), and no main effect of treatment (F(1,48)= 0.13, p>0.05). Similar the optical stimulation findings, a two-way ANOVA on median amplitude (Fig. 5B_2_) revealed a significant interaction (F(1,48)= 10.90, p<0.01), a main effect of exposure (F(1,48)= 5.01, p<0.05), and a main effect of treatment (F(1,48)= 15.42, p<0.001). Bonferroni posttests indicated a significant increase in the aEPSC amplitude of saline injected CIE/WD animals (11.1 ± 0.6pA, N=11) compared to saline injected AIR-exposed controls (9.0 ± 0.6pA, N=12; p<0.01, t=3.44), to neurons from ketamine injected Air-exposed animals (8.7±0.2pA, N=13; p<0.01, t=4.02), and to CIE/WD neurons from animals injected with ketamine (8.2 ± 0.3pA, N=16; p<0.001, t=5.13). Notably, aEPSC amplitudes recorded from neurons within the CIE/WD + ketamine group were not significantly different neurons recorded from either the saline injection Air (p>0.05, t=1.50) or ketamine injected Air groups (p>0.05, t=0.96). These data suggest that ketamine’s effect on CIE/WD aESPC amplitude is generalizable to the population of EC-BLA synapses.

### Ketamine administration at the onset of withdrawal attenuates withdrawal-induced anxiety-like behavior

The reversal of CIE/WD-dependent postsynaptic facilitation of NMDA and AMPA receptor function suggested that ketamine may also have striking effects on withdrawal-related behaviors. To examine this, we assessed the effects of ketamine administration on anxiety-like behavior on the elevated zero maze 24h into withdrawal following 10 days of CIE (Fig. 6C). A two-way ANOVA of time spent in open areas on the EZM (Fig. 6A) found a significant interaction (Exposure X Treatment; F(1,32)= 11.70, p<0.01) but no main effects of treatment (Saline vs Ketamine; F(1,32)= 0.15, p>0.05) or exposure (Air vs CIE/WD; F(1,32)=0.86, p>0.05). Multiple comparison posttests indicated that saline injected CIE/WD animals spent significantly less time in the open areas of the EZM (30.7 ± 8.0sec, N=9) compared to saline injected AIR-exposed control animals (92.2 ± 21.7sec, p<0.01, t=3.08, N=9). CIE/WD animals injected with ketamine spent significantly more time (73.7 ± 14.0sec, p<0.05, t=2.15, N=9) in the open areas than CIE/WD animals injected with saline. Surprisingly, AIR animals injected with ketamine spent significantly (p < 0.01) less time (38.6 ± 8.2sec, p<0.05, t=2.69, N=9) in the open areas than AIR animals injected with saline. Air-ketamine and CIE-saline groups were not significantly different from each other (p>0.05, t=0.39). These findings paralleled observations with respect to head-dips (explorations over the side of the open portions of the apparatus) which also showed a significant interaction between exposure (AIR vs CIE/WD) and treatment (saline vs ketamine; F(1,32)=9.55, p<0.01) along with significant posttest differences between Air-saline (70.6±5.3) and CIE/WD-saline (41.7±8.1, p<0.01, t=2.93), CIE/WD-saline and CIE/WD-ketamine (63.8±6.7, p<0.05, t=2.24), and Air-saline and Air-ketamine groups (49.6±7.5, p<0.05, t=2.12). Similarly, there was a significant interaction between the main factors for closed ➔ open transitions (F(1,32)=8.03, p<0.01) which produced significant differences between Air-saline (6.1±1.3) and CIE/WD-saline groups (2.2±0.5, p<0.01, t=2.75) and Air-saline and Air ketamine groups (3.2±0.8, p<0.05, t=2.04) along with a trend for differences between the CIE/WD-saline and CIE/WD-ketamine groups (5.0±1.2, p∼0.06, t=1.96). A two-way ANOVA of total distance moved (Fig. 6B) found no main effect of exposure (F(1,31)= 1.322, p= 0.2591), no main effect of treatment (F(1.31)= 0.09139, p= 0.7644), and no interaction (F(1,31)= 1.569, p= 0.2198), indicating that ketamine did not have any significant effects on locomotion. These data suggest that a single ketamine injection delivered at the end of the last CIE exposure and ameliorate withdrawal-related anxiety-like behavior measured 24h later.

**Figure 6.**
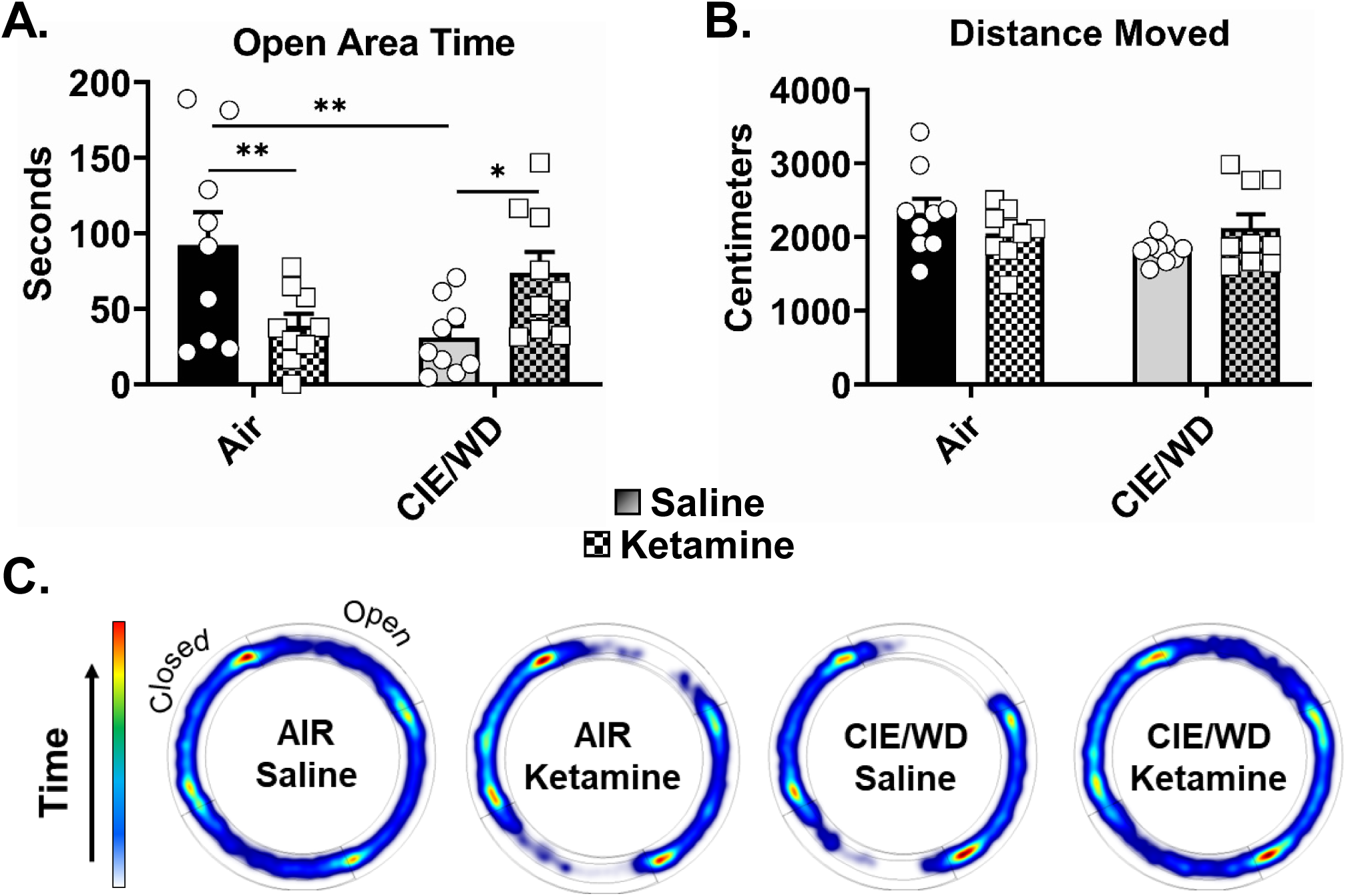
Ketamine administration at the onset of withdrawal prevents ethanol withdrawal-associated increases in anxiety-like behavior. **A**., Time spent in the open areas of the EZM measured 24 hours after air or 10-days of CIE ±ketamine administration (10mg/kg, IP). Animals in the CIE/WD + Saline group (N= 9) spent significantly less time (Two-way ANOVA, interaction p<0.01, F(1,32)=11.7) in the open areas compared to the AIR + Saline (N= 9, Bonferonni posttest, ** - p<0.01) and CIE/WD + Ketamine groups (N= 9, * - p<0.05). AIR + Ketamine animals (N= 9) spent significantly less time in the open compared to Air + Saline (** - p<0.01). **B**, There were significant differences in total distance moved between any of the treatment groups (Two-way ANOVA, interaction p>0.05, F(1,32)=4.0; exposure p>0.05, F(1,32)=0.15; treatment p>0.05, F(1,32)=0.90). **C**, EZM heat plots representing group data for times spent in the various parts of the apparatus.

## Discussion

In this study we examined the effects of chronic ethanol exposure and withdrawal on glutamate transmission in the AIC – BLA circuit. Using a combination of optogenetics and electrophysiology, we demonstrate that neurons in the AIC make monosynaptic glutamatergic synapses with principal neurons in the BLA (Fig. 1). Following withdrawal from 10 days (but not 7 days) of CIE, AIC – BLA synapses show increased postsynaptic function demonstrated by an increase in the amplitude of AMAPR-mediated aEPSCs recorded using strontium substitution (Fig. 3). Notably, neither 7 or 10 days of CIE produced presynaptic changes in glutamate release probability from AIC – BLA synapses (Fig. 2). Importantly, a subanesthetic dose of ketamine at the onset of withdrawal prevented increases in postsynaptic function at AIC – BLA synapses. Further, this ‘protective’ effect was generalizable to the population of EC synapses activated by electrical stimulation (Fig. 5). We also found that ketamine prevented ethanol-induced changes in NMDA receptor function by using electrical stimulation to record an NMDAR-mediated input-output curve for EC – BLA synapses (Fig. 4). Finally, we examined withdrawal-associated anxiety-like behavior using the EZM and found that ketamine attenuates the development of increased negative-affect in CIE/WD animals (Fig. 6). Surprisingly, we found that ketamine alone produced an opposite effect in air-exposed control animals, increasing anxiety-like behavior relative to saline-injected controls. Together, these results indicate that postsynaptic function of the AIC – BLA pathway is sensitive to chronic ethanol exposure and that acute ketamine is an effective modulator of both this glutamate plasticity in the BLA and anxiety-like behavior.

The effects of chronic ethanol on the AIC – BLA pathway had not been examined prior to this study. However, previous studies demonstrate that both the AIC and the BLA are independently altered by chronic ethanol exposure. For example, chronic ethanol self-administration in monkeys resulted in greater glutamatergic EPSCs within the AIC as compared to the ethanol-naïve group [35]. The insular cortex in alcohol dependent human subjects exhibits greater atrophy compared to the non-dependent controls; and, impaired cognitive functioning in these alcohol users [36] suggests the insular cortex is particularly vulnerable to chronic ethanol exposure. In the amygdala, alcohol dependence affects a broad range of systems including synaptic transmission, neurotransmitter transport, structural plasticity, metabolism, energy production, transcription and RNA processing, and the circadian cycle [37]. Our laboratory has reported that chronic ethanol exposure increases presynaptic glutamate release probability and synaptic glutamate concentrations and decreases synaptic ‘failures’ at *stria terminalis* inputs using electrical stimulation [31]. Additionally, we’ve reported that chronic ethanol increases postsynaptic NMDAR function measured from ‘local’ BLA synapses [28], increases synaptic responses mediated by both kainate [29]- and AMPA-type [10] glutamate receptors at EC – BLA inputs. Our current findings demonstrating that AIC – BLA synapses express post- but not presynaptic plasticity are in line with published work showing that ethanol-induced changes in glutamate synaptic function are input-dependent. Given the increased amygdala and insula reactivity to certain types of emotional processing [38], our findings help support the notion that chronic ethanol disrupts emotional-processing across a range of contributing neural circuits and neurophysiological mechanisms.

With electrical stimulation, we’ve previously shown that postsynaptic changes at EC-BLA synapses occur following 7 days of CIE exposure in adolescent (∼P35) animals. But, in the present study, 10 days CIE, but not days, increased in postsynaptic function in AIC – BLA synapses using optogenetic activation of these inputs. While it is possible that the AIC-BLA synapses are more resilient to chronic ethanol compared to the population of EC inputs in which they reside, a more parsimonious explanation for the relative delay in the development of CIE-dependent postsynaptic facilitation is the age of the animals during the ethanol exposure.

Expression of channelrhodopsin in the AIC terminal fields requires several weeks; and, as a result, young adults (∼P80) were subjected to the CIE. Indeed, others have also reported that adolescent animals are more vulnerable to ethanol (reviewed in [39]), highlighting an interesting age-depend decrease in ethanol sensitivity, even at the level of the synapse.

NMDA receptor-dependent synaptic plasticity, such as long-term potentiation (LTP) and long-term depression (LTD), has received much attention given its role in learning and memory. The initial expression of this type of plasticity is mediated by a redistribution of AMPA-type glutamate receptors and, with time, structural changes which require the synthesis of new proteins. Strong evidence suggests that postsynaptic potentiation is a result of increases in calcium influx via the opening of NMDARs, which can activate calcium/calmodulin-dependent kinase II (CamKII). This leads to phosphorylation of a number of proteins including AMPARs, which can cause an increase in the conductance of the AMPAR channel and the insertion of AMPARs into the postsynaptic membrane [40]. In line with this, we have reported that chronic ethanol exposure increases CamKII activation and phosphorylation [10], which may help explain the increases in AMAPR receptor function measured with electrophysiology and suggests that ethanol-induced plasticity may occur through a similar mechanisms as NMDAR-dependent LTP. In the present study, we tested this hypothesis by injecting animals with a subanesthetic dose of ketamine, a NMDAR antagonist, at the onset of withdrawal would alter glutamate plasticity in withdrawal. We saw profound effects of ketamine on both NMDAR and AMPAR function.

Specifically, ketamine blocked the ethanol-induced increase in NMDAR function (Fig. 4), which we hypothesize may be responsible for the attenuation of the ethanol-induced increase in AMPAR function (Fig. 5). Additionally, we found that ketamine was able to prevent ethanol withdrawal-induced increase in anxiety-like behavior. Our findings parallel those of a recent study that found ketamine administered at the onset of forced abstinence from chronic two-bottle choice ethanol drinking prevented abstinence-dependent anxiety on the elevated plus maze and restored the capacity for LTP within the bed nucleus of the stria terminalis (BNST) which is normally occluded by chronic ethanol [34]. Interestingly, Vranjkovic and colleagues report that ketamine administered either two- or 6-days post-abstinence failed to prevent these abstinence-induced changes, suggesting the timing of administration is critical. This appears consistent with our own findings that increased AMPA receptor synaptic function at EC synapses occurs within the first 24h of withdrawal from CIE [10]. Notably, ketamine administration in air-exposed control animals resulted in a significant increase in anxiety-like behavior (Fig. 6) without altering glutamate function in the AIC – BLA or EC – BLA pathway (Fig. 5). First, this suggests to us that the anxiogenic effect of ketamine in control animals is not mediated by changes in BLA glutamate transmission. Interestingly, others have also reported anxiogenic effects of ketamine [41–43], although these effects are usually at slightly higher doses than what we used and sometimes after repeated administration. One of these studies found that atypical antipsychotic drugs such as clozapine and risperidone counteract ketamine-induced alterations in a variety of anxiety-related behaviors [41], suggesting that our findings could potentially be explained by ketamine acting on serotonin and dopamine systems.

Second, we find it interesting that ketamine alone is more ‘chronic ethanol’-like with respect to behavioral outcomes. Notably, ethanol and ketamine both inhibit NMDAR function suggesting ketamine’s behavioral effect in air-exposed animals could therefore be related to the consequences of this interaction. Regardless, our findings are consistent with the notion that ketamine may be protective against chronic ethanol-induced changes in glutamate synaptic plasticity and negative affective disturbances at least in the short-term.

Both clinical and preclinical studies suggest that a subanesthetic-dose of ketamine serves as a fast-acting antidepressant (reviewed in [44]). And, the U.S. Food and Drug Administration recently approved esketamine, the s-enantiomer of ketamine, nasal spray for treatment-resistant depression [45]. However, the efficacy of ketamine in the treatment of substance use disorders, including AUD, is less characterized. A recent systematic review of clinical AUD studies found improvements in abstinence rates for up to two years following a single infusion of ketamine [46]. There are also two ongoing clinical trials evaluating the use of ketamine in the treatment of AUD, one looking at ketamine alone and in combination with naltrexone for the rapid treatment of MDD and AUD (NCT02461927) and another examining ketamine for reduction of alcoholic relapse (NCT02649231). Collectively, these studies suggest that ketamine could improve abstinence in AUD.

In summary, we demonstrate that AIC – BLA synapses express increases in postsynaptic AMPA receptor function following withdrawal from 10 days of CIE. We hypothesized that this may be a result of increased NMDA receptor function at EC – BLA synapses. Both of these functional alterations are blocked by a subanesthetic dose of ketamine administration at the onset of ethanol withdrawal. Finally, acute ketamine administration at the onset of withdrawal was able to attenuate anxiety-like behavior. Future studies will focus on established a role of AIC – BLA circuit in anxiety-like behavior as well as trying to further elucidate the mechanisms by which ketamine produces these outcomes.

